# Plant traits are poor predictors of long-term ecosystem functioning

**DOI:** 10.1101/859314

**Authors:** Fons van der Plas, Thomas Schröder-Georgi, Alexandra Weigelt, Kathryn Barry, Sebastian Meyer, Adriana Alzate, Romain L. Barnard, Nina Buchmann, Hans de Kroon, Anne Ebeling, Nico Eisenhauer, Christof Engels, Markus Fischer, Gerd Gleixner, Anke Hildebrandt, Eva Koller-France, Sophia Leimer, Alexandru Milcu, Liesje Mommer, Pascal A. Niklaus, Yvonne Oelmann, Christiane Roscher, Christoph Scherber, Michael Scherer-Lorenzen, Stefan Scheu, Bernhard Schmid, Ernst-Detlef Schulze, Vicky Temperton, Teja Tscharntke, Winfried Voigt, Wolfgang Weisser, Wolfgang Wilcke, Christian Wirth

## Abstract

Earth is home to over 350,000 vascular plant species^1^ that differ in their traits in innumerable ways. Yet, a handful of functional traits can help explaining major differences among species in photosynthetic rate, growth rate, reproductive output and other aspects of plant performance^2–6^. A key challenge, coined “the Holy Grail” in ecology, is to upscale this understanding in order to predict how natural or anthropogenically driven changes in the identity and diversity of co-occurring plant species drive the functioning of ecosystems^7, 8^. Here, we analyze the extent to which 42 different ecosystem functions can be predicted by 41 plant traits in 78 experimentally manipulated grassland plots over 10 years. Despite the unprecedented number of traits analyzed, the average percentage of variation in ecosystem functioning that they jointly explained was only moderate (32.6%) within individual years, and even much lower (12.7%) across years. Most other studies linking ecosystem functioning to plant traits analyzed no more than six traits, and when including either only six random or the six most frequently studied traits in our analysis, the average percentage of explained variation in across-year ecosystem functioning dropped to 4.8%. Furthermore, different ecosystem functions were driven by different traits, with on average only 12.2% overlap in significant predictors. Thus, we did not find evidence for the existence of a small set of key traits able to explain variation in multiple ecosystem functions across years. Our results therefore suggest that there are strong limits in the extent to which we can predict the long-term functional consequences of the ongoing, rapid changes in the composition and diversity of plant communities that humanity is currently facing.

## BODY

Worldwide, ecological communities are rapidly changing due to various anthropogenic activities^9–12^. This biodiversity change is non-random, and the functional traits of organisms driving their growth, survival and reproduction are key in determining which species thrive and which perish under global change^13–15^. This may have important implications, as traits not only affect individual plant performance, but they may also drive various ecosystem functions such as biomass production, and the services these functions provide to human well-being^7, 8, 15^.

Predicting rates of ecosystem functioning, such as biomass production or carbon sequestration, from the composition or diversity of traits in plant communities has been coined the “Holy Grail” in ecology^7, 8^. Various studies have shown links between plant traits and *species-level* variation in photosynthetic rate, growth, and reproductive output present in the plant kingdom^3–5^. However, in natural communities, plants occur in various abiotic environments, and they interact with individuals from other species, so that both the identity and diversity of traits may matter for *ecosystem-level* functioning. Despite this, so far various field studies only found relatively weak links between the identity and diversity of plant traits and ecosystem-level functioning^8, 16–18^. Furthermore, those studies that did find strong links between traits and ecosystem functions^19, 20^ were typically carried out within a single year, but if links between traits and ecosystem functioning are highly context-dependent, the capacity of traits to predict the long-term consequences of global change, thereby attaining the “Holy Grail”, may still be limited. Alternatively, strong and consistent links between plant traits and ecosystem functioning exist, but higher numbers and more appropriate traits than assessed in previous studies are needed to demonstrate those links.

To test these ideas, we first performed a systematic literature review to investigate which and how many traits 100 recent studies measured when attempting to link the diversity or composition of traits within terrestrial plant communities to ecosystem functioning. We found that most studies analyzed six traits, and only two studies assessed more than 15 traits (Fig. 1B). Nine of the ten most frequently studied traits (Fig. 1A) described aboveground plant properties, of which six described leaf properties. Only one frequently measured trait was related to plant roots, even though roots provide important plant functions (e.g. anchoring, resource uptake) and represent approximately 50% of total plant biomass^21^. Thus, most previous studies assessed a sparse set of traits, with a strong bias towards leaf traits.

**Figure 1.**
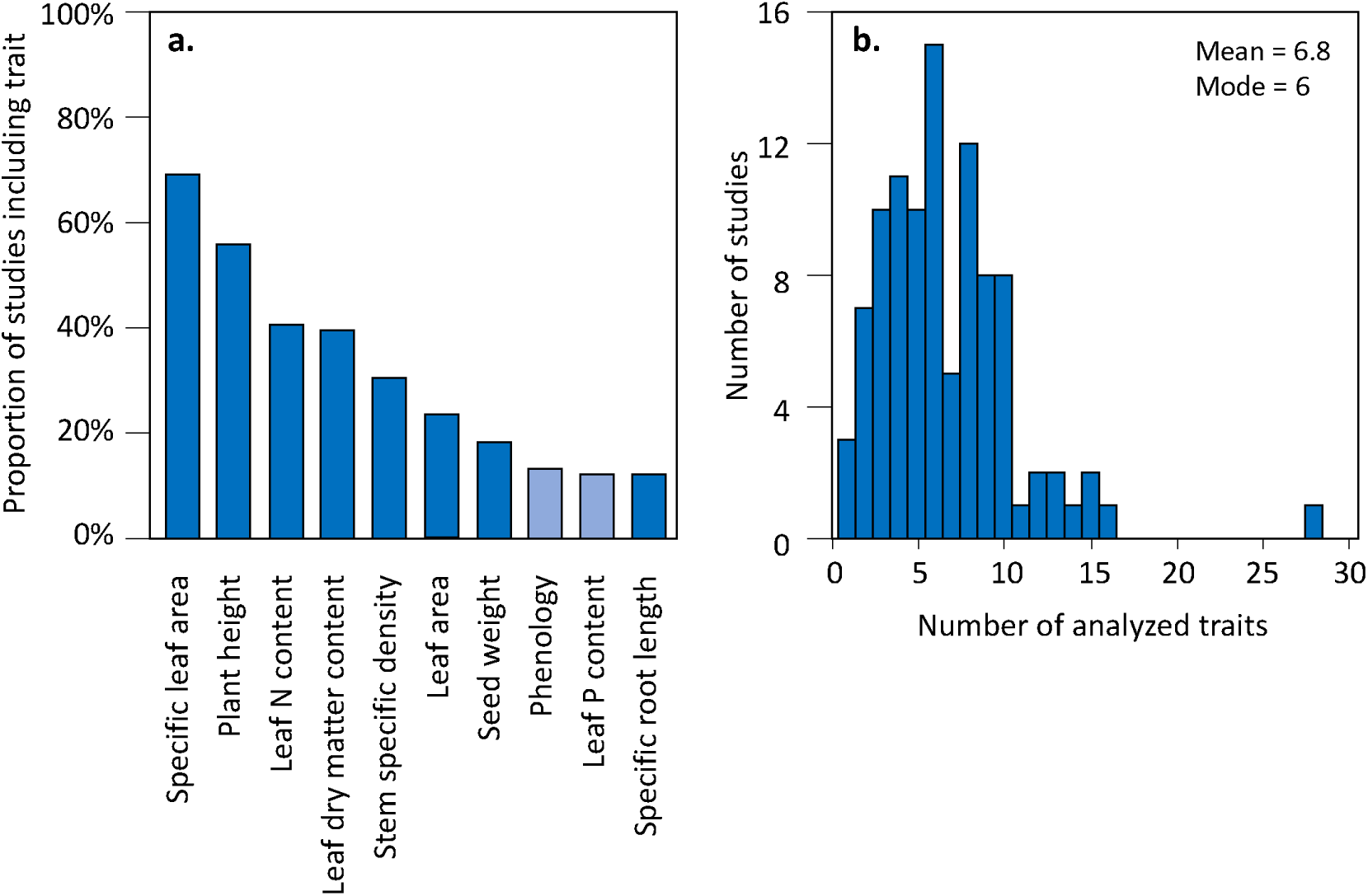
Overview of which and how many traits are typically analyzed in other ecosystem functioning-related studies. A: Percentage of studies in which the 10 most frequently measured traits were investigated, according to the review of 100 recently published articles. The lighter blue bar shows the only two functions not measured in this study. B: Number of measured traits among studies.

We then investigated to what extent a much higher number of traits can explain variation in ecosystem functioning. We did this using a dataset containing 10 years of measurements of 42 ecosystem functions, assessed in 78 experimentally established grassland communities in Germany. The 42 ecosystem functions described various above- and belowground stocks and rates of plant, faunal, and abiotic properties driving grassland functioning (Supplementary Methods). Both the diversity and composition of the studied plant communities were experimentally manipulated, by sowing different combinations of species^22, 23^. For each species, we measured 41 traits (more than any of the studies assessed in our review) related to structural, morphological, chemical and physiological properties of all main plant parts, including leaves, stems, flowers, seeds, and roots. By combining these trait data with plant community data, we quantified both the Functional Identity and the Functional Diversity for each plot in each year. Functional Identity was calculated as the abundance-weighted mean of a trait within a community, and drives ecosystem functioning if the contributions of species to ecosystem functioning are proportional to their relative abundance^15, 24^. Functional Diversity was calculated as Rao’s Quadratic Entropy^25^, and can drive ecosystem functioning if species contribute differently to functioning when co-occurring with plants with different traits, e.g. due to trait-driven resource complementarity^23, 25, 26^.

We used linear mixed models to analyze how much of the variation of each of the 42 ecosystem functions was explained by Functional Identity and Diversity metrics of all 41 traits, as well as by random year and plot differences. We used a forward model selection procedure, in which during each step a trait was added, if it significantly improved model fit and did not strongly correlate with the traits already present in the model. Despite the high number of traits included in our analysis, and even though each ecosystem function was on average driven by 4.8 traits (Fig. 2B), the average marginal R^2^ of final models was 0.127, indicating that traits explained on average only 12.7% (ranging from 0.0% to 40.0%) of the variation in ecosystem functioning (Fig. 2C). Marginal R^2^ values were even lower (mean of 0.078) when we used a more conservative model selection procedure correcting for False Discovery Rates. Conditional R^2^ values, which also account for the variance explained by random factors, including year differences, were much higher, with an average value of 0.632. Our finding that traits explained a very low proportion of variance may seem surprising, as other studies explained more variance with fewer predictors^19^. However, other studies typically used data for single years only, and it is possible that links between traits and ecosystem functions are only strong within years. To test this, we also analyzed links between ecosystem functions and traits for each year separately. This showed that within years marginal R^2^ values were much higher, with an average value of 0.326. Thus, while traits were poorly linked to ecosystem functioning across years (possibly due to strong shifts in species’ abundances^75^), they explained much more variation within years, indicating that links between traits and ecosystem functions are strongly context-dependent.

**Figure 2.**
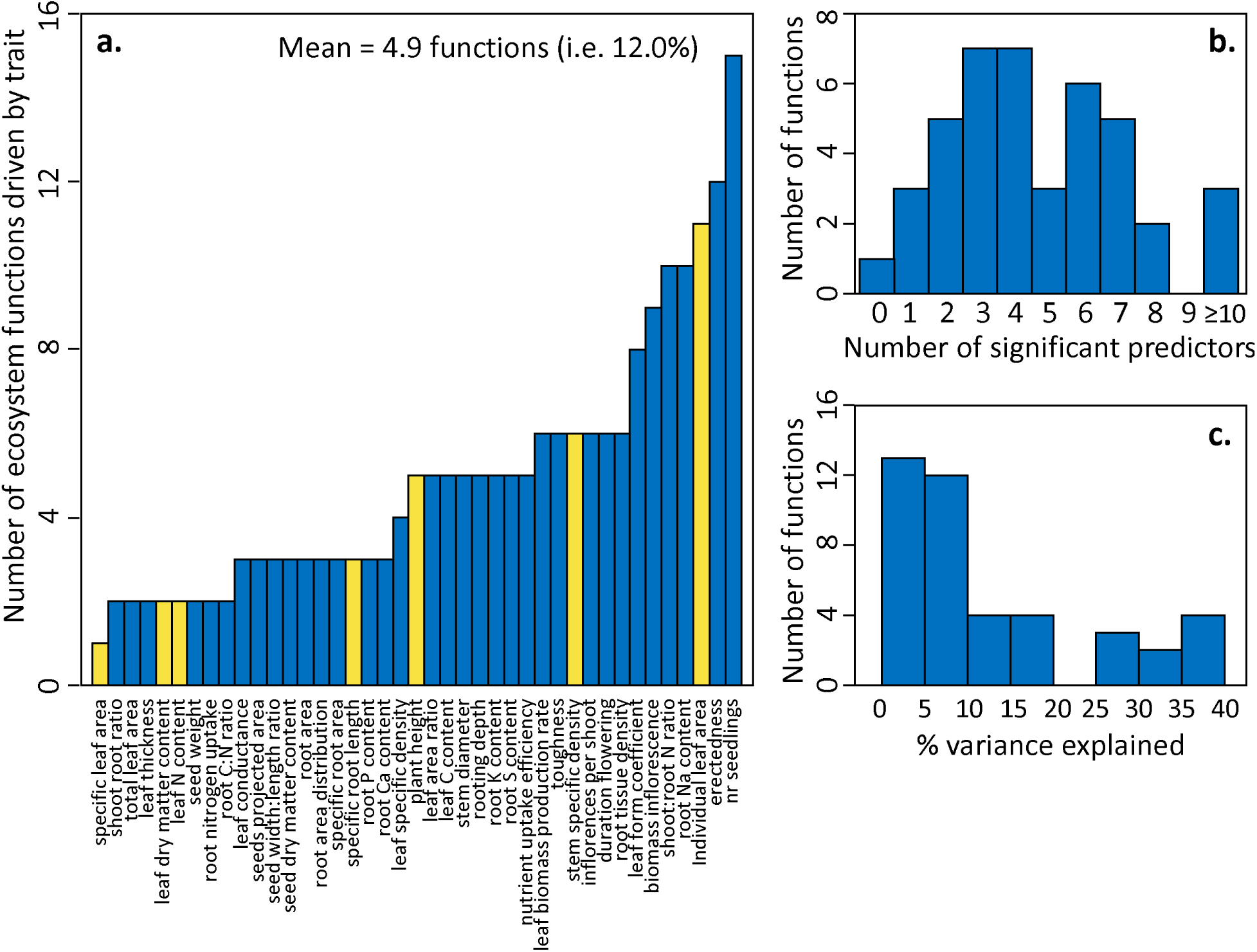
The relative importance of different and multiple traits for ecosystem functioning across years. A: the number of analyzed functions that was significantly driven by each trait, according to final models. The traits analyzed in over 10% of the papers included in the review are shown in yellow. B: Number of significant predictors in final models of each ecosystem function. C: Marginal R^2^ values for final models of each ecosystem function.

We then assessed how our ability to explain rates of ecosystem functions across years depends on how many and which traits are included in analyses. Those traits most frequently assessed in other studies did not drive more functions than traits less frequently studied. One trait (specific leaf area) only significantly drove a single ecosystem function, while others (e.g. leaf area) drove many more, but an overall pattern was not detectable (Fig. 2A). We investigated more formally how our ability to explain variation in ecosystem functioning would change, if we had measured either *a*) a random subset of six (corresponding to the number of traits assessed in most other studies) out of the 41 traits (based on 100 random draws), or *b*) only the six traits most frequently assessed in other studies, or if *c*) we analysed species richness (the most commonly used biodiversity indicator) instead as a predictor of ecosystem functioning. Irrespective of whether six random traits or those most frequently investigated in other studies were analysed, on average only 4.8% (95 percentile: 3.8-6.5%) of ecosystem functioning variation could be explained (Fig. 3A,B), while species richness could explain only 1.7% of variation in ecosystem functioning. This represents a strong decrease compared to the 12.7% of variation explained when all 41 traits were assessed (Fig. 2B). We also assessed to which extent analyzing subsets of fewer or more than six traits influenced the proportion of explained variance in ecosystem functioning. This showed that there was an asymptotic relationship between the number of traits analyzed and the average proportion of explained variation in ecosystem functioning, and that at least 9, or 24 traits are required to explain 5%, and 10% of the variation in ecosystem functioning, respectively (Fig. 4A).

**Figure 3.**
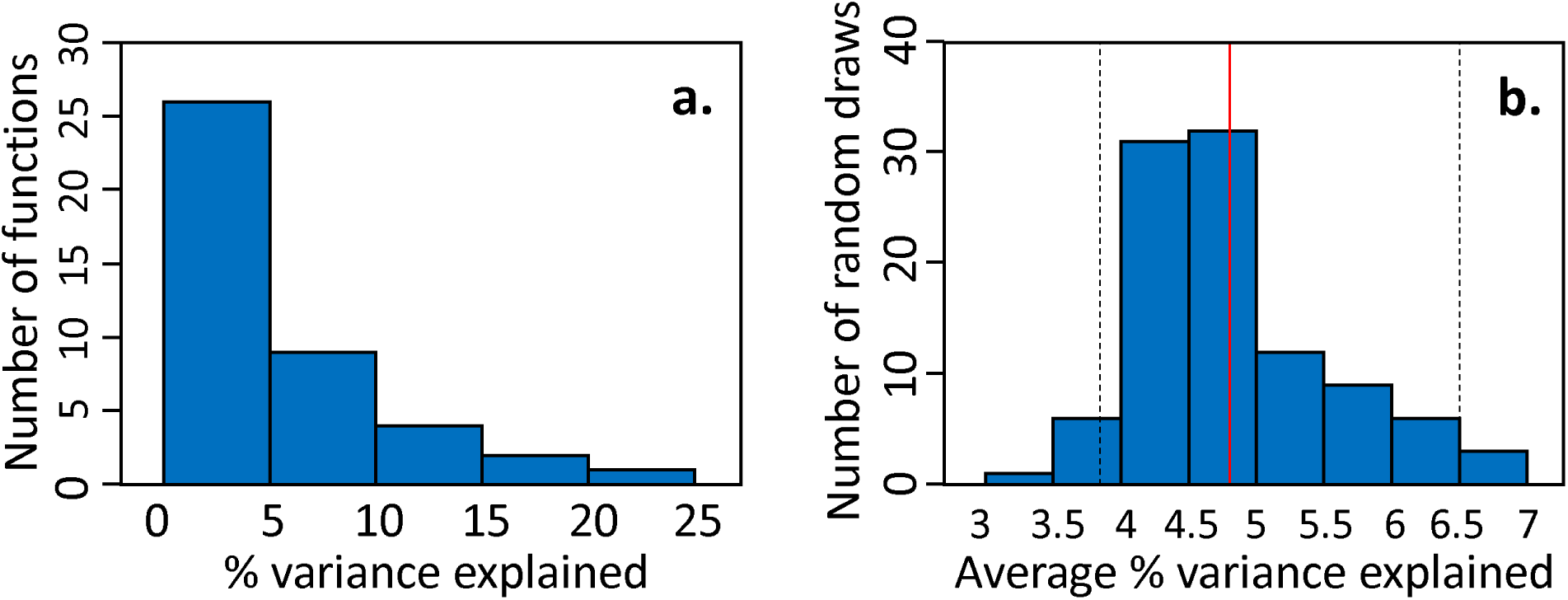
R^2^ values of models in which only six traits were analyzed to explain ecosystem functions across years. A: Distribution of marginal R^2^ values of final models for each trait, when only the six most frequently investigated traits (see review) were included in the analysis. B: Distribution of mean marginal R^2^ values (across final models for each trait), when based on 100 random draws, six randomly selected investigated traits were included in the analysis. The vertical dashed bars show the 95% confidence interval, while the vertical red bar shows the mean marginal R^2^ across all functions when only the six most frequently investigated traits were included in the analysis.

**Figure 4.**
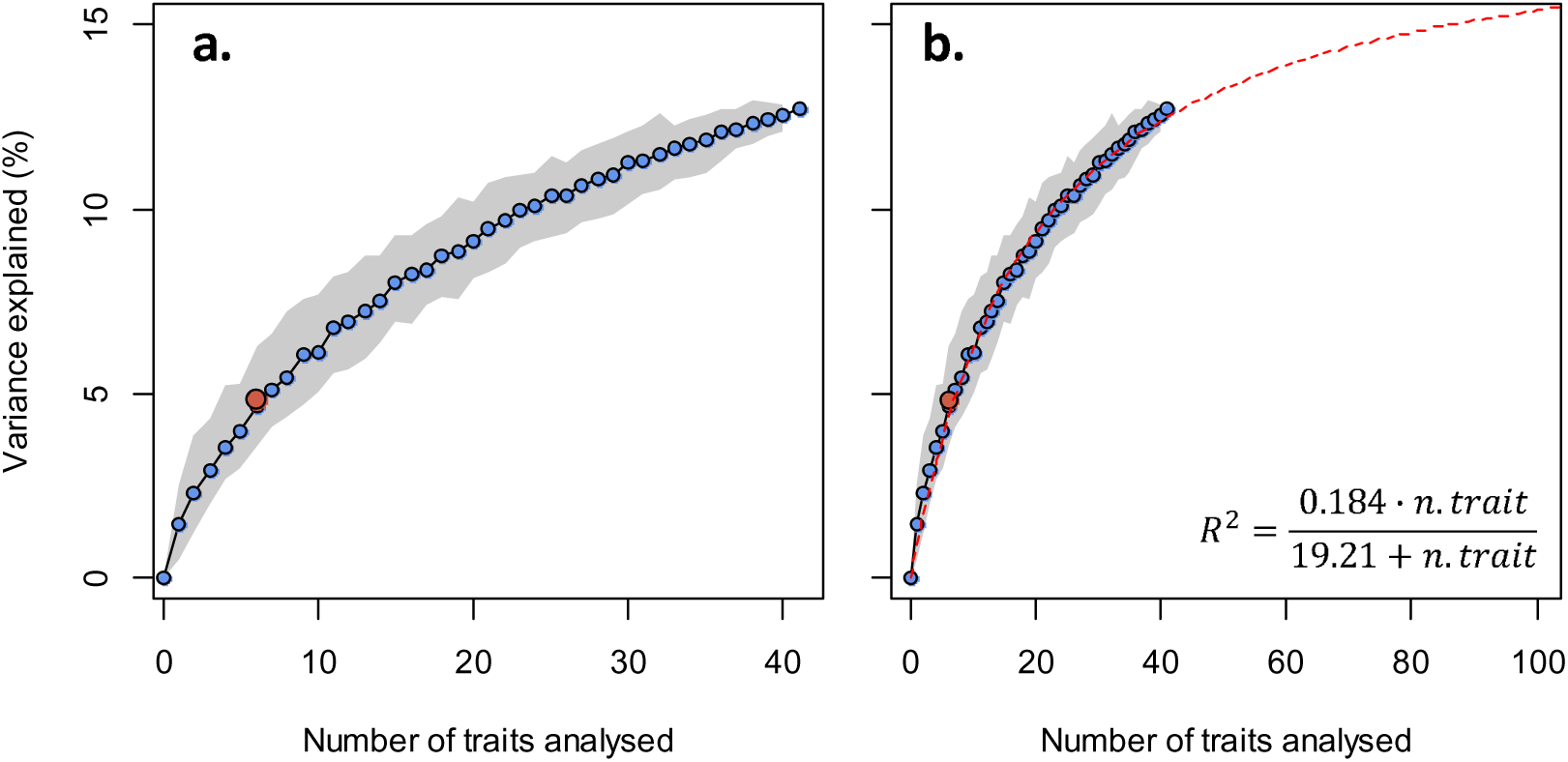
The average proportion of variation in ecosystem functions across years explained by plant traits increases asymptotically with the number of traits included in the analysis. The red dot shows the proportion of explained variation when only the six traits most commonly assessed in other studies are included. A: the marginal R^2^ – number of traits relationship based on analysis of actual data. B: an additional extrapolated (based on a fitted Michaelis – Menten equation) marginal R^2^ – number of traits relationship (red, dashed line).

Thus, while each ecosystem function alone was on average explained by fewer than 5 traits (Fig. 2B), many more traits are needed to explain multiple ecosystem functions (Fig. 4). While seemingly a paradox, this happens if different ecosystem functions are driven by different traits. We demonstrated this by calculating the overlap (*o*) in the traits significantly driving each pair of ecosystem functions, using Sørenson’s index^27^. The average overlap indicated that pairs of ecosystem functions had on average only 12.2% significant trait drivers in common. Thus, while traits are commonly advertised as conveying more general information than a species’ identity does^7, 14, 26^, a small set of key traits able to explain variation in multiple ecosystem functions does not exist in Central European grasslands, just like ‘superspecies’ providing multiple functions don’t exist^28^.

While many ecosystem functions were relatively poorly explained by traits, we could nevertheless identify traits that predicted many ecosystem functions, and ecosystem functions that were better predicted by traits than others. All traits explained at least one ecosystem function, and some (e.g. leaf area) drove many more (Fig. 2A). We also found that ecosystem functions related to aboveground stocks or processes were much better predicted (average marginal R^2^ = 0.21) than those related to belowground stocks or processes (average marginal R^2^ = 0.07) (Table S2.1), even though 14 root traits were included in our analysis. It is possible that unmeasured traits related to litter quality or mycorrhizal associations have stronger links to functions such as soil respiration or soil nutrient availability. However, extrapolation of the observed relationships between model *R^2^* and the number of analysed traits suggests that 87 traits are needed to increase the proportion of variance explained to 15%, and that there is an upper limit of around 18% in the proportion of variance explained, even if an unlimited number of traits is analyzed (Fig. 4B). Hence, the inclusion of more trait data would only yield limited gains in our ability to explain ecosystem functioning. Instead, it is possible that the inclusion of intraspecific variation (not considered in this study) would improve links with ecosystem functions^29^. In addition, there were small spatial mismatches between within-plot locations of ecosystem function measurements and vegetation surveys, which could have weakened links between traits and ecosystem functioning. Lastly, it is possible that traits are more strongly linked to ecosystem functioning within other systems such as forests, or across ecosystem types.

Using one of the most comprehensive studies so far, we showed that while traits can be strongly linked to ecosystem functions within years, our capacity to predict levels of multiple ecosystem functions across years (differing in e.g. weather conditions) is strongly limited. Thus, finding ecology’s Holy Grail is extremely challenging at best, and at worst a ‘mission impossible’. This may have strong implications. The functional composition and diversity of plant communities are rapidly changing^9–12^, and researchers are employing increasingly complex models to predict the consequences of these changes for worldwide biogeochemical and hydrological cycles^30, 31^. While we encourage the use of such models and their inclusion of increasingly accurate trait information, our work also raises concerns about limits in their predictive capacity, suggesting that the consequences of ongoing biodiversity change are largely unpredictable. Human well-being relies on ecosystem services that are underpinned by various ecosystem functions^32, 33^, and insuring that these functions are provided at high levels is extremely challenging if future environments are dominated by plant communities differing from those observed today. Hence, policies halting the current-day, rapid changes in biodiversity are the safest bet to guarantee nature’s contributions to future generations of people.

## ACKNOWLEDGEMENTS

We thank Enrica de Luca, Anja Vogel, Helmut Hillebrand and Elisabeth Marquard for their contributions to the data collection. The Jena Experiment is funded by the German Science Foundation (DFG Oe516/3-1, 3-2, 10-1).

## AUTHOR CONTRIBUTIONS

F.v.d.P., T.S-G., A.W., K.B. and C.W. conceived the ideas and designed the study. F.v.d.P., T.S-G., S.M. and A.A. performed the analyses. All authors, except for F.v.d.P., K.B. and A.A., contributed to the data collection. F.v.d.P wrote a first draft of the paper, and all other authors contributed to editing several manuscript versions.

## METHODS

### Review

We performed a review to investigate which traits were most often analyzed as predictors of ecosystem functioning in recent years. We did this on the Clarivate Analytics Web of Science website in July 2018, using the search terms (functional-diversity *or* community-weighted-mean *or* CWM *or* trait-diversit*) *and* ecosystem function* *and* (plant *or* vegetation). This initially yielded 654 results. Among these, we searched for papers that analyzed an ecosystem function (broadly defined as energy or trophic fluxes and biomass stocks, measured at the ecosystem or community level) as the response of the Functional Diversity or Functional Identity (e.g. (abundance-weighted) trait mean values) of one or more terrestrial plant traits. We only focused on the 100 most recently published articles that met these criteria. The main objective of this mini-review was to get an overview of a representative sample of recent studies linking terrestrial plant traits to ecosystem functioning, rather than to get an exhaustive overview of all published literature.

Among the 100 selected papers (see Appendix A), we screened which plant traits were analyzed as predictors of ecosystem functioning. Some traits had different labels among different publications (e.g. specific leaf area versus leaf mass per area^34, 35^. In those cases, we used our expert judgement and a plant trait thesaurus (http://www.top-thesaurus.org/home)^36^ to relabel traits in order to obtain a common terminology. We then counted and ranked the frequencies (number of papers) by which each trait was analyzed as a predictor of ecosystem functioning, and we identified the top ten of traits analyzed in most papers, and the five most commonly analyzed traits.

### Experimental design

We studied relationships between various ecosystem functions and plant traits using data from the Jena Main Biodiversity Experiment^22, 23^, which is one of the biggest and longest running biodiversity experiments worldwide. This grassland biodiversity experiment was set up in spring 2002 in the floodplain of the Saale river close to the city of Jena (Germany, 50°55’N, 11°35’E, 130 m a.s.l.), at a field that was previously managed as a fertilized agricultural field for at least four decades. The experiment was designed to study the effects of species and functional group richness on various ecosystem functions.

In short, 78 plots were established, each measuring 20×20 m. In these plots, different subsets of a species pool of 60 species were sown in spring 2002. The different species were selected to be representative of a Molinio-Arrhenatheretea grasslands^37^ and were classified in four functional groups as ‘grass’ (including Poaceae and one Juncaceae species), small herb, tall herb or legume, with 16, 12, 20 and 12 species in the species pool, respectively. In each plot, 1, 2, 4, 8 or 16 species were sown, with each richness level replicated 16 times. The 16 species mixture plots formed an exception, and were replicated only 14 times. Total sowing density was 1000 seeds per m^2^, irrespective of the richness level. Each plot contained a unique species composition. In addition to a species richness gradient, a functional group richness gradient was established, in such a way that sown species and functional group richness were as orthogonal as possible. Functional group richness ranged from 1, 2, 3 and 4, with 34, 20, 12 and 12 replicates, respectively. Plots were assigned to four blocks in parallel to the riverside to account for differences in soil properties with increasing distance from the river (with e.g. sand content being higher in plots closer to the Saale river). Each block had a similar number of plots, and each block had all levels of species and functional group richness approximately equally represented.

Twice per growing season, plots were weeded in order to avoid species that were not sown in the plots upon establishment. We refer to two other publications^22, 23^ for more details on the design of the Jena main experiment.

### Plant community assessments

During the period between 2003 and 2012, twice per year, during spring (May) and summer (August), cover of all target plant species was estimated in each plot, within a 3×3 m subplot. For more details, we refer to Roscher et al. (2013)^38^.

### Ecosystem function measurements

During the years 2002 till 2012, 42 different ecosystem variables (‘ecosystem functions’ hereafter) were measured, describing plant, faunal and abiotic pools and process rates, some of which were measured aboveground, and some of which were measured belowground. Some ecosystem functions were measured in multiple seasons or years, always using standardized protocols. The ecosystem functions measured were: plant biomass consumed by herbivores, herbivory rate, frequency of pollinator visits, abundance of soil surface fauna, richness of soil surface fauna, abundance of vegetation layer fauna, richness of vegetation layer fauna, number of pollinator species, drought resilience, drought resistance, leaf area index, bare ground cover, aboveground plant biomass, dead plant biomass, cover of invasive plant species, richness of invasive plant species, rain throughfall, basal soil respiration, soil respiratory quotient, earthworm biomass, soil larvae abundance, soil mesofauna abundance, soil macrofauna abundance, biomass of soil microbes, biomass of plant roots, downward flux water in upper soil, downward flux water in deeper soil, upward flux water in upper soil, upward flux water in deeper soil, evapotranspiration in upper soil, evapotranspiration in deeper soil, upper soil water content, deep soil water content, inorganic carbon content, organic carbon content, soil bulk density, soil nitrogen content, soil δ^15^N values, soil NH_4_ content, soil NO_3_ content, nitrate leaching and soil phosphorus content (see Table S1.1 for a more detailed overview). When ecosystem functions were measured multiple times within a year (e.g. both in spring and summer) within the same plot, we used averages of those repeated measurements in further analyses. For detailed descriptions on the methodology of all ecosystem function measurements, we refer to the Supplementary Materials.

### Trait measurements

In total, 41 plant traits were measured. These traits described whole plant, leaf, stem, flower, seed, (fine) root characteristics, and were structural, morphological, chemical, physiological, phenological. The measured traits included all terrestrial plant traits identified as ‘most commonly assessed’ in our mini-review, except for leaf phosphorus content. For a complete overview of all measured traits, we refer to Table S1.2. The majority of the traits, including most leaf and root traits, were measured in mesocosms filled with Jena field soil mixed with sand in the Botanical Garden of Leipzig (Saxony, Germany), in 2011 and 2012. Mass fraction and number of inflorescences and seedling density were measured in monocultures at the Jena Experiment. Rooting depth and flower duration could not be reliably estimated in the 80 cm high mesocosms and was therefore derived from earlier published measurements^20^. Detailed information on the individual trait measurements is provided in Supplementary Material.

### Quantifying Functional Diversity and Functional Identity

We combined the species-level abundance assessments for each plot with the trait measurements to quantify Functional Diversity and Identity in each plot, separately for each combination of year and season. Functional Diversity was calculated for each trait (thus yielding 42 Functional Diversity measures in total) separately using Rao’s Quadratic Entropy metric^24^ (or Q), which measures the sum of pairwise trait distances of co-occurring species, whereby pairwise distances are weighted by the relative abundance of the species: 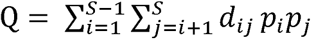, where *i* and *j* are the two species forming a species pair, *S* is the species richness within a community, *d_ij_* is the Euclidean trait distance and *p_i_* and *p_j_* are the relative abundance of species *i* and *j*, respectively. Here, relative abundances are measured as the species’ cover (estimated in subplots of 3 x 3 m, see above) within a plot divided by the total community cover. Functional Identity was measured for each trait (thus also yielding 41 measures in total) using the Community Weighted Mean (CWM) metric^15^, which measures the abundance-weighted average of trait values among species within a community as: 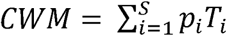, where *T_i_*. indicates the trait value of species *i*. We also recalculated FD and CWMs based on presence-absence data (thus ignoring differences in relative abundance of species present in a plot) for sensitivity analyses.

In addition to calculating CWM and FD values, we also calculated the realized species richness for each plot and each year, based on the species-level abundance assessments.

### Statistical analyses

We first analyzed how each ecosystem function was related to all 41 measured traits. This was done using a separate Linear Mixed Model (LMM) for each function, in which the CWM and Rao’s Q values for each trait were treated as fixed factors (thus yielding 2 × 41 = 82 fixed factors), and year and plot were treated as random factors. We used a forward model selection procedure, in which first ‘empty’ models only containing random factors were fitted, and then significant fixed factors were added step-by-step. We chose a forward model selection procedure to overcome problems related to multicollinearity (many traits, and hence FD and FI metrics, were correlated, see Table S2.2). During each step in our selection procedure, we first tested for the significance of all *n* fixed factors (where *n* = the total number of 82 fixed factors minus the number of fixed factors already included at earlier steps of the model selection procedure) that could be added to the previous, less complex model, using log-likelihood tests. We then investigated which factor was most significant, and added this factor to the previous model if it did not lead to any Variance Inflation Factor (VIF) exceeding 5. In case the most significant fixed factor did cause multicollinearity (maximum VIF > 5), we investigated if the next-most significant factor could be added. This procedure was repeated until we ended up with a model only containing significant fixed factors with VIF values ≤ 5, to which no significant (P ≤ 0.05) fixed factors could be added. LMM fitting was done using a Restricted Maximum Likelihood procedure, using the lmer function of the lme4 package^39^ in R-3.5.1^40^. We calculated the marginal (proportion of variance exclusively explained by fixed factors, i.e. traits) and conditional (proportion of variance explained by fixed factors and random factors combined) R^2^ values^41^ using the r.squaredGLMM function of the MuMIn package^42^ in R-3.5.1^40^. We also performed some sensitivity analyses, in which we repeated the above analyses, with *i*) as the only difference that we corrected for False Discovery Rates^43^, to reduce the risk of type I errors, *ii)* as the only difference that FD and CWM values based on presence-absence data were used as predictors and *iii*) where we replaced FD and CWM predictor variables by realized species richness.

We also investigated to which extent links between the Functional Diversity and Identity of traits and ecosystem functions changed, if we analysed ecosystem functions for each year in which they were measured separately. We did this by running the same models and model selection procedure as described above, except that the random factor ‘year’ was omitted from the models (as functions were analyzed for each year separately, this random factor had become obsolete). In addition, the random factor ‘plot’ was omitted from the models, as we only had one measurement per plot within each year.

To quantify the overlap in significant predictors among different ecosystem functions, we created a 42 (number of ecosystem functions) × 41 (number of traits) binary matrix, with cells containing values of 1 when either the FD and/or the FI of the corresponding trait significantly drove the ecosystem function, and a value of 0 when neither the FD nor the FI significantly drove the ecosystem function. We then calculated the overlap (*o*) in the sets of traits significantly driving each pair of ecosystem functions, using Sørenson’s index^28^ as: 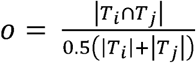 where |*T_i_*| and |*T_i_*| are the numbers of traits significantly driving respectively ecosystem function *i* and *j*, and |*T_i_* ∩ *T_j_*| is the number of traits significantly driving both ecosystem function *i* and *j* and we then calculated the average overlap. Importantly, these overlap estimates could be conservative (i.e. underestimated) due to strong correlations between traits. Therefore, we repeated the above described linear mixed models (originally with 82 fixed factors, corresponding to the FD and FI values of 41 traits), but then using Principal Component Analysis (PCA) axis values based on the FD and FI values as explanatory variables. To this end, we first performed a PCA, and we selected the 13 PCA axes that explained more than 100/82 (the number of input variables) = 1.22% of all FD and FI variation. Together, these 13 PCA axes explained 92% of all FD and FI variation. The selection procedure of models linking ecosystem functions with PCA axes was the same as for the main analyses linking ecosystem functions with FD and FI variables. We then repeated the overlap analysis in the same way as described above, and found that for FD and FI metrics based on PCA variables, the average overlap of 25.7% was somewhat, but not much, higher than the overlap based on FD and FI metrics of raw traits.

We then analyzed to what extent a subset of the six traits most commonly assessed in other studies, i.e. specific leaf area, plant height, leaf N concentration, leaf dry matter content, stem tissue density and leaf area, could explain variance in ecosystem functioning. To this end, we repeated the modeling procedure described above, except that only the above mentioned six traits were assessed in the model selection procedure, rather than the full set of 41 traits. In addition, we also assessed how random subsets of *n* traits, with n ranging from 1 to 40, could explain ecosystem functioning. To this end, we ran 100 simulations for each level of *n*. In each of these simulations, we first randomly selected a subset of *n* traits out of the total of 41 traits. For these random subsets of *n* traits, we again ran the same model selection procedure as described above for each ecosystem function, to assess which of the traits significantly drove the levels of each function, and in order to assess the marginal R^2^ values of final models. For each simulation, we then calculated the mean (across all functions) marginal R^2^ value, and for each *n*, we calculated the mode and 95% percentiles for the mean marginal R^2^ value across the 100 simulations (as reported in Fig. 4). Only for *n* = 1 and *n* = 40 traits this procedure was slightly different, as for both of these levels of *n*, there were only 41 traits or trait combinations possible. Thus, in those cases, we did not take 100 random draws of traits, but instead systematically analysed at all possible combinations. Based on the resulting relationship between the number of traits analyzed and the marginal R^2^ values, we fitted a non-linear model using the nls function in R3.5.3, of the form: 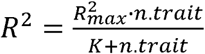 in which R^2^ is the marginal R^2^ value, 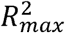 is the asymptote in marginal R^2^ value, n.trait the number of traits analysed, and K describes the slope by which the 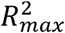 is reached. The resulting 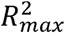 and *K* values were 0.184 and 19.21 respectively, and these were used to extrapolate the observed relationship between the number of traits analyzed and the marginal R^2^ values, in order to calculate how many traits were required to obtain marginal R^2^ values of 0.150 and higher.

## SUPPLEMENTARY MATERIALS

### S1. SUPPLEMENTARY METHODS

#### S1.1. Ecosystem function measurements

During the years 2002 until 2012, 42 different ecosystem functions were measured. Some ecosystem functions were measured in multiple seasons or years, although always using standardized protocols. An overview of the different ecosystem functions can be seen in Table S1.1.

**Table S1.1.** List of all ecosystem functions analyzed in this study.

| Ecosystem function | unit | Summary description | Years measured |
| --- | --- | --- | --- |
| Consumed plant biomass | $\text{g m}^{-2}$ | Biomass consumed by herbivores | 2010-2012 |
| Herbivory rate | % | % of leaves damaged | 2003-2005, 2010-2012 |
| Frequency pollinator visits | nr | Number of observed pollinator visits | 2005, 2006, 2008 |
| Abundance soil surface fauna | nr | Abundance of invertebrates caught in pitfall traps | 2003, 2005, 2010 |
| Richness soil surface fauna | nr | Species richness of invertebrates caught in pitfall traps | 2003, 2005, 2010 |
| Abundance vegetation layer fauna | nr | Abundance of invertebrates caught via suction sampling | 2003, 2005, 2010 |
| Richness vegetation layer fauna | nr | Species richness of invertebrates caught via suction sampling | 2003, 2005, 2010 |
| Number of pollinator species | nr | Number of observed pollinator species | 2005, 2006, 2008 |
| Drought resilience | $\text{g m}^{-2}$ | Resistance biomass production after drought | 2009-2012 |
| Drought resistance | $\text{g m}^{-2}$ | Resistance biomass production to drought | 2008-2012 |
| Leaf Area Index | unitless | Leaf area index (measure of light interception) | 2003-2012 |
| Bare ground cover | % | Cover of bare ground | 2002-2011 |
| Target plant biomass | $\text{g m}^{-2}$ | Aboveground dry mass of target species | 2002-2012 |
| Dead plant biomass | $\text{g m}^{-2}$ | Aboveground dry mass of dead target species | 2003-2008 |
| Cover invasive species | % | Cover of non-target plant species | 2003-2007 |
| Richness invasive species | nr | Number of non-target plant species | 2003-2007 |
| Rain throughfall | mm | Amount of rainwater reaching lower vegetation layers | 2008-2012 |
| Basal soil respiration | $\mu\text{L g}^{-1} \text{h}^{-1}$ | Basal soil respiration (proxy of decomposition) | 2003-2008, 2010-2012 |
| Soil respiratory quotient | $\mu\text{L g}^{-1} \text{h}^{-1}$ | Respiration per biomass soil microbes | 2008, 2010-2012 |
| Earthworm biomass | g | Biomass of earthworms | 2003-2008 |
| Soil larvae abundance | nr | Number of larvae in soil | 2004, 2006, 2008 |
| Soil mesofauna abundance | nr | Count of mesofauna individuals in soil | 2004, 2006, 2008 |
| Soil macrofauna abundance | nr | Count of macrofauna individuals in soil | 2004, 2006, 2008 |
| Biomass soil microbes | $\mu\text{g C g}^{-1}$ | Biomass of microbes in soil | 2003, 2004, 2006-2008, 2010-2012 |
| Biomass plant roots | g | Belowground plant biomass in soil | 2003, 2004, 2006-2008, 2011 |
| Downward flux water upper soil | $\text{L m}^{-2}$ | Downward flux of water in upper soil | 2003-2007 |
| Downward flux water deep soil | $\text{L m}^{-2}$ | Downward flux of water in deeper soil | 2003-2007 |
| Upward flux water upper soil | $\text{L m}^{-2}$ | Upward flux of water in upper soil | 2003-2007 |
| Upward flux water deep soil | $\text{L m}^{-2}$ | Upward flux of water in deeper soil | 2003-2007 |
| Evapotranspiration upper soil | $\text{L m}^{-2}$ | Evapotranspiration in upper soil | 2003-2007 |
| Evapotranspiration deep soil | $\text{L m}^{-2}$ | Evapotranspiration in deeper soil | 2003-2007 |
| Upper soil water content | $\text{L m}^{-2}$ | Water content in upper soil | 2003-2007 |
| Deep soil water content | $\text{L m}^{-2}$ | Water content in deeper soil | 2003-2007 |
| Inorganic soil carbon | % | Concentration of inorganic carbon in soil | 2002, 2004, 2006 |
| Organic soil carbon | % | Concentration of organic carbon in soil | 2002, 2004, 2006 |
| Bulk density soil | $\text{g m}^{-3}$ | Bulk density soil (proxy for compaction) | 2002, 2004, 2006 |
| Nitrogen content soil | % | Soil total nitrogen content | 2002, 2004, 2006 |
| Soil $\delta^{15}\text{N}$ | ‰ | Soil nitrogen isotope ratios | 2002, 2004, 2006 |
| Soil $\text{NH}_4$ content | $\mu\text{g g}^{-1}$ | Soil ammonium concentration | 2002-2008 |
| Soil $\text{NO}_3$ content | $\mu\text{g g}^{-1}$ | Soil nitrate concentration | 2002-2008 |
| Nitrate leaching | $\text{mg m}^{-2}$ | Nitrate leaching | 2002-2006 |
| Soil phosphate content | $\text{mg L}^{-1}$ | Soil phosphate content | 2003-2007, 2009, 2011, 2012 |

##### S1.1.1. Consumed plant biomass

Herbivory rates were converted into estimates of consumed plant biomass in three steps. First, the total leaf biomass of a species in a plot was estimated from the species-specific aboveground biomass that included the biomass of leaves, stems, and inflorescences, using the ratio of leaf biomass to total aboveground biomass. Second, the leaf biomass of each species in each mixture was multiplied by the respective herbivory rate to obtain the leaf biomass consumed from this species in gram dry weight per square meter. Third, the total biomass removed from a particular plant community was calculated by summing the consumed leaf biomass over all plant species in the community^44, 45^.

##### S1.1.2. Herbivory rate

Large vertebrates were excluded from the experimental site by a fence such that herbivory was only caused by invertebrates (though there was occasional grazing by voles). Herbivory was measured during the biomass harvest twice a year – typically at the end of May and the end of August. Herbivory was measured in five years (2012 to 2014)^44, 45^. For each target species present in the sorted biomass samples, usually, 30 fully developed leaves (only 20 in 2012 and 2013) were sampled randomly for herbivory measurements. For species with fewer than the target number of leaves in the sample, all available leaves were measured. The leaf area of all sampled leaves (i.e. the area left after feeding of the herbivores including petioles) was measured with a leaf area meter (LI-3000C Area Meter, LI-COR Biosciences, Lincoln (NE), USA). Herbivore damage (i.e., the leaf area damaged by herbivores in mm^2^) was estimated visually by comparing the damaged leaf area to a series of circular and square templates ranging in size from 1 mm^2^ to 500 mm^2^. Herbivory damage included four different herbivory damage types: chewing, sap sucking, leaf mining and rasping damage. For each leaf, a single value of the total area damaged by all types of herbivory was estimated. Herbivory rates (the proportion of leaf area damage) for each plant species in a mixture was calculated by dividing the estimated area damaged by herbivores by the original leaf area without damage. To obtain the total leaf area before herbivore feeding, we summed the leaf area remaining after feeding by herbivores that was measured with a leaf-area meter and the leaf area removed by chewing herbivores using plant species-specific ratios of herbivory damage types. A community level herbivory rate was calculated by summing the species-specific herbivory rates weighted by their respective relative leaf biomass for each biomass sample. For a detailed description of the methodology used see Meyer et al. 2017^45^.

##### S1.1.3. Frequency of pollinator visits

We observed flower-pollinator interactions within a quadrat of 80×80cm three times during the vegetation period in 2005, 2006 and 2008^46, 47^. During the six-minute observation period every interaction was counted as a flower visitation. Observations were only conducted on sunny days between 09:00 and 17:00 h.

##### S1.1.4. Fauna soil surface abundance

For recording the activity abundance of ground-dwelling arthropods, we installed two pitfall traps of 4.5 cm diameter per plot in 2003, 2005, and 2010^48, 49^. Traps were replaced six times in 2003 and 2005 between May and October, and every two weeks between May and September in 2010. In the field we filled traps with 3% formalin, and stored them later in 70% ethanol.

##### S1.1.5. Fauna soil surface species richness

For recording the activity abundance of ground-dwelling arthropods, we installed two pitfall traps of 4.5 cm diameter per plot in 2003, 2005, and 2010^48, 49^. Traps were replaced six times in 2003 and 2005 between May and October, and every two weeks between May and September in 2010. In the field we filled traps with 3% formalin, and stored them later in 70% ethanol.

##### S1.1.6. Fauna vegetation abundance

For recording the abundance of vegetation-associated arthropods we used suction sampling in 2003, 2005, 2010^48, 49^. Five (2003 and 2005) and nine (2010) times during the vegetation period we randomly placed cages of 0.75 m3, cleared them from arthropods, and stored all sampled animals in 70% ethanol.

##### S1.1.7. Fauna vegetation species richness

For recording the species richness of vegetation-associated arthropods we used suction sampling in 2003, 2005, 2010^48, 49^. Five (2003 and 2005) and nine (2010) times during the vegetation period, we randomly placed cages of 0.75 m3 and cleared them from arthropods. We stored all sampled animals in 70% ethanol and sent them to external taxonomists for species-level identification.

##### S1.1.8. Pollinator species richness

We observed flower-pollinator interactions within a quadrat of 80×80cm three times per year in 2005, 2006 and 2008^46, 47^. During the six-minute observation period we identified every flower-visiting insects to species or morphospecies. Unknown species were captured for later identification. Observations were only conducted on sunny days between 09:00 and 17:00 h.

##### S1.1.9. Drought resilience

We used data from the drought experiment established as 1×1 m subplots on 76 plots of the Jena Main Experiment in 2008. The two subplots per plot were designated as either drought or ambient control using rainout shelters constructed using wooden frames and transparent PVC roofs^50^ (see Vogel et al. 2013 for details). Rainwater was collected in rain barrels and used to water ambient subplots following rainfall events^50, 51^. Shelters were set up mid-summer and excluded natural rainfall from mid-July to the end of August (six weeks). Standing biomass was harvested in May and August (before removal of the shelters) as described for standing aboveground biomass.

We calculated resilience from our biomass data according to van Ruijven and Berendse^52^. Resilience determines the change in biomass production after perturbation and was calculated as difference of post-drought biomass and the corresponding ambient treatment from the first harvest after drought (May the following year).

##### S1.1.10. Drought resistance

Drought resistance was calculated based on the same data as drought resilience (S1.1.9). We calculated resistance from our biomass data according to van Ruijven and Berendse^52^ as the difference of biomass under perturbed and unperturbed conditions (drought - ambient) at the end of the drought period in August.

##### S1.1.11. Leaf area index

Community leaf area index (LAI) was measured twice a year just before biomass harvest (see S1.1.13) with a LAI-2000 plant canopy analyzer (LI-COR) using high resolution and a view cap masking 45° of the azimuth towards the operator. In 2003 and 2004, 10 randomly allocated measurements were taken at 5 cm height within an area of 3 x 3 m in the center of the core area. From 2005 onwards all measurements were taken along a 10 m transect in the core area of each experimental plot. One above reading was taken at the first transect point, followed by 10 below readings taken with 1 m distance from each other. We used the mean over the 10 calculated LAI values from the below readings as mean community LAI per plot.

##### S1.1.12. Bare ground cover

Bare ground cover was visually estimated together with sown species cover in September 2002 and twice a year just before biomass harvest. Bare ground cover was estimated directly as percentage of area. From 2002 to 2004, measurements were taken in two extra carefully weeded sub-areas of 2 x 2.25 m. We report the average value based on these two estimates for community cover. From 2005 onwards all measurements were taken in one 3 x 3 m area in the core area of each experimental plot.

##### S1.1.13. Target aboveground plant biomass

Aboveground community biomass was harvested twice a year just prior to mowing (during peak standing biomass in late May and in late August) on all experimental plots. This was done by clipping the vegetation at 3 cm above ground in two to four randomly selected rectangles of 0.2 x 0.5 m per plot. The harvested biomass was sorted into sown species, total weeds and detached dead organic material and dried to constant weight (70°C, ≥ 48 h). Target aboveground plant biomass was calculated as the sum of biomass for all sown species from all rectangles per plot.

##### S1.1.14. Dead plant biomass

Sum of biomass of detached dead organic material from all rectangles per plot as described in target aboveground plant biomass.

##### S1.1.15. Cover invasive species

Cover of invader species was visually estimated to the nearest percentage before weeding (spring = April, summer = July) on the same subplot size as used for the quantification of invader species richness (S1.1.16) in each large plot from 2003 to 2007. In the field, invader species cover was separately recorded for internal invader species (i.e. species belonging to the experimental species pool, but not to the sown species composition of the respective plot) and external invader species (i.e. species not belonging to the experimental species pool). Cover of internal and external invader species was summed to get the total cover of invader species^53^.

##### S1.1.16. Richness invasive species

Within each large plot one subplot of 2.00 × 2.25 m was permanently marked to quantify invasion resistance from 2003 to 2007. All invader species present in this subplot were recorded before weeding (spring = April, summer = July) to assess invader species richness^53^.

##### S1.1.17. Rain throughfall

In biweekly intervals from 2008 to 2012, throughfall volume was collected with rain collectors (2-L sampling bottles connected to funnels [diameter of 0.12 m], both polyethylene). The sampling bottles were protected against larger particles and small animals with a polyethylene net (0.005 m mesh width). The collectors were cleaned with deionized water before installation and replaced by clean collectors in 2- to 3-month intervals.

##### S1.1.19. Basal soil respiration

In each year, five randomly located soil samples were taken per plot with a soil corer (5 cm diameter, 5 cm deep) and pooled plot-wise. Before measuring, all samples were homogenized, sieved (2 mm), larger roots and soil animals were picked by hand, and samples were stored in plastic bags at 5°C. Microbial respiration was measured using an electrolytic O_2_-microcompensation apparatus^54^. O_2_ consumption of soil microorganisms in ∼5 g of fresh soil (equivalent to c. 3.5 g soil dry weight) was measured at 22°C over a period of 24 h. Basal respiration [µL O_2_ g^-1^ dry soil h^-1^] was calculated as mean of the O_2_ consumption rates of hours 14 to 24 after the start of the measurements.

##### S1.1.19. Soil respiratory quotient

In each year, five randomly located soil samples were taken per plot with a soil corer (5 cm diameter, 5 cm deep) and pooled plot-wise. Before measuring, all samples were homogenized, sieved (2 mm), larger roots and soil animals were picked by hand, and samples were stored in plastic bags at 5°C. Microbial respiration was measured using an electrolytic O_2_-microcompensation apparatus^54^. O_2_ consumption of soil microorganisms in ∼5 g of fresh soil (equivalent to c. 3.5 g soil dry weight) was measured at 22°C over a period of 24 h. Basal respiration [µL O_2_ g^-1^ dry soil h^-1^] was calculated as mean of the O_2_ consumption rates of hours 14 to 24 after the start of the measurements. Substrate-induced respiration (SIR) was determined by adding D-glucose to saturate catabolic enzymes of the microorganisms according to preliminary studies (4 mg D-glucose g^-1^ dry soil solved in 400 µL deionized water^55^. The maximum initial respiratory response (MIRR; [µL O_2_ g^-1^ dry soil h^-1^]) was calculated as mean of the lowest three O_2_-consumption values within the first 10 h after glucose addition. Microbial biomass carbon [µg C g^-1^ dry soil] was calculated as 38 × MIRR^56^. The soil respiratory quotient was calculated by dividing basal respiration by microbial biomass^57^.

##### S1.1.20. Earthworm biomass

Earthworm extractions were performed on one subplot of 1 x 1 m per plot that was established to extract earthworms repeatedly. Subplots were enclosed with PVC shields aboveground (20 cm) and belowground (15 cm). Two earthworm extraction campaigns were performed twice per year in spring and autumn of 2005, 2006, and 2008 by electro-shocking^58^. Therefore, a combination of four octet devices (DEKA 4000, Deka Gerä tebau, Marsberg, Germany; Thielemann^59^) was used which were powered by two 12 V car batteries. Eight steel rods (length 60 cm) were inserted into the soil (to a depth of w55 cm) per octet device forming four circles of six rods (each 50 cm in diameter) with two rods in the center of each circle. An electrical voltage was applied in pulses to the moist soil (earthworm extractions were always performed during humid and mild weather conditions) sequentially to pairs of rods in the circle (negative pole) and in the center of the circle (positive pole). In each subplot earthworm extraction was performed for 35 min, increasing the voltage from 250 V (10 min) to 300 V (5 min), 400 V (5 min), 500 V (5 min), and 600 V (10 min). Despite the PVC shields, earthworms re-colonized earthworm subplots until the next extraction campaign^58^. Extracted earthworms were identified, counted and weighed in the laboratory.

##### S1.1.21. Soil larvae abundance

Soil macrofauna was collected from soil cores taken to a depth of 10 cm in autumn 2004 (October), 2006 (November) and 2008 (October). Soil cores were taken using a steel corer (22 cm diameter). One soil core per plot was taken, and soil animals were extracted by heat^60^, collected in diluted glycerol, and transferred into ethanol (70%) for storage. Soil animals were identified^61–63^ and counted. A detailed list of soil animal taxa and their trophic assignment is given in Eisenhauer et al. (2011)^64^.

##### S1.1.22. Soil mesofauna abundance

Soil mesofauna was collected from soil cores taken to a depth of 10 cm in autumn 2004 (October), 2006 (November) and 2008 (October). Soil cores were taken using a steel corer (5 cm diameter). One soil core per plot was taken, and soil animals were extracted by heat^60^, collected in diluted glycerol, and transferred into ethanol (70%) for storage. Soil animals were identified^65–67^ and counted. A detailed list of soil animal taxa and their trophic assignment is given in Eisenhauer et al. (2011)^64^.

##### S1.1.23. Soil macrofauna abundance

Soil macrofauna was collected from soil cores taken to a depth of 10 cm in autumn 2004 (October), 2006 (November) and 2008 (October). Soil cores were taken using a steel corer (22 cm diameter). One soil core per plot was taken, and soil animals were extracted by heat^60^, collected in diluted glycerol, and transferred into ethanol (70%) for storage. Soil animals were identified^65–67^ and counted. A detailed list of soil animal taxa and their trophic assignment is given in Eisenhauer et al. (2011)^64^.

##### S1.1.24. Soil microbial biomass

In each year, five randomly located soil samples were taken per plot with a soil corer (5 cm diameter, 5 cm deep) and pooled plot-wise. Before measuring, all samples were homogenized, sieved (2 mm), larger roots and soil animals were picked by hand, and samples were stored in plastic bags at 5°C. Soil microbial biomass respiration was measured using an electrolytic O_2_-microcompensation apparatus^54^. O_2_ consumption of soil microorganisms in ∼5 g of fresh soil (equivalent to c. 3.5 g soil dry weight) was measured at 22°C over a period of 24 h. Substrate-induced respiration (SIR) was determined by adding D-glucose to saturate catabolic enzymes of the microorganisms according to preliminary studies (4 mg D-glucose g^-1^ dry soil solved in 400 µL deionized water^55^). The maximum initial respiratory response (MIRR; [µL O_2_ g^-1^ dry soil h^-1^]) was calculated as mean of the lowest three O_2_-consumption values within the first 10 h after glucose addition. Microbial biomass carbon [µg C g^-1^ dry soil] was calculated as 38 × MIRR^56^. The soil respiratory quotient was calculated by dividing basal respiration by microbial biomass^57^.

##### S1.1.25. Plant root biomass

Standing root biomass was sampled down to 30 cm depth in all plots in June 2003, September 2004, and June 2006, 2008 and 2011. Two monoculture plots were excluded because of poor establishment. In all years we took several soil cores per plot and processed the pooled samples (2003: 5 cores with 4.8 cm diameter; 2004: 3 cores with 4.8 cm diameter; 2006: 5 cores with 8.7 cm diameter; 2008: 3 cores with 4.8 cm diameter; 2011: 3 cores with 3.5 cm diameter). The cores were cooled (4 °C; frozen in 2006) until further handling. The bulk material of the pooled cores was weighed and cut to 1 cm pieces before subsampling. For root washing, a 50 g subsample was soaked in water and then repeatedly rinsed with tap water over a 0.5 mm sieve. In 2011, the full bulk sample was washed for root material. Roots were dried at 60 – 70 °C and weighed subsequently.

##### S1.1.26. Upper (0-30 cm) and deep (0-70 cm) soil water content

Volumetric soil water contents were measured with frequency domain reflectometry (FDR) using a mobile manual FDR probe (PR1/6 and PR2/6, Delta-T-Devices, Cambridge, UK) on all plots in 1–2 weekly resolution in the 0.1, 0.2, 0.3, 0.4, and 0.6 m soil depths^68, 69^.

Soil water contents per plot were aggregated to depth-weighted means for the 0-0.3 m (“upper soil”) and 0.3-0.7 m (“deep soil”) soil layers. At a central automatic meteorological station on the field site, soil water contents in the 0.08, 0.16, 0.32, and 0.64 m soil depths were measured with Theta Probe soil moisture sensors – ML2x (Delta-T Devices, Cambridge, UK) in 10-min resolution between 1 July 2002 and 31 December 2007 and aggregated to daily depth-weighted means for the 0.0-0.3 and 0.3-0.7 m soil layers. To obtain a complete soil water contents data set for the 0.0-0.3 and 0.3-0.7 m soil layer per plot for the years 2003-2007, data gaps were filled with Bayesian hierarchical models using the soil water contents from the central meteorological station as explanatory variable^70^.

##### S1.1.27. Downward and upward flux and evapotranspiration of soil water, in upper and deep soil

A water balance model was used to simulate downward and upward water fluxes and actual evapotranspiration from the 0-0.3 m (“upper soil”) and the 0.3-0.7 m (“deep soil”) soil layers per plot for the years 2003-2007 in weekly resolution^70^. The model uses the input variables precipitation (measured at the central meteorological station in 10-min resolution), potential evapotranspiration (calculated from meteorological data from the central station using the Penman-Wendling equation), and volumetric soil water contents (see S1.1.26). The model is based on the water balance equation: precipitation + upward flux = downward flux + actual evapotranspiration - change in volumetric soil water content between two subsequent observation dates. The percentage of roots in each soil layer was used as a proxy for the percentage of potential evapotranspiration that could be evaporated from the respective soil layer. Together with using the net flux (downward flux - upward flux) from the upper soil layer as input into the deep soil layer, this allowed for modeling of the water fluxes for the two soil layers 0-0.3 m and 0.3-0.7 m separately^70^.

##### S1.1.28. Inorganic and organic soil carbon

Total carbon concentration was analyzed biannually on ball-milled sub-samples by an elemental analyzer at 1150 °C (Elementaranalysator vario Max CN, Elementar Analysensysteme GmbH, Hanau, Germany). To determine the organic carbon concentration we measured inorganic carbon concentration by elemental analysis at 1150 °C after removal of organic carbon for 16 h at 450 °C in a muffle furnace. Organic carbon concentration was then calculated from the difference between both measurements^71, 72^.

##### S1.1.29. Soil bulk density

In 2002, soil bulk density in the plough horizon was determined on 27 plots from undisturbed soil samples with a depth resolution of 10 cm. The respective samples were taken with a metal bulk density ring of 10 cm height, passed through a sieve with 2 mm mesh size, dried to constant weight at 105 °C and were subsequently weighed to calculate the density. The chosen plots represented a spatial gradient across the field site and resulted in average soil bulk density estimations at the beginning of the experiment. Starting in 2004 all bi-annually soil samples were taken with the split tube sampler, dried and weighed to detect changes in the bulk density. The inner diameter of the soil corer was used for volume calculation^71^.

##### S1.1.30. Total soil nitrogen

Total nitrogen concentration was analyzed bi annually on ball-milled sub-samples by an elemental analyzer at 1150 °C (Elementaranalysator vario Max CN, Elementar Analysensysteme GmbH, Hanau, Germany)^71, 72^.

##### S1.1.31#Soil δ ^15^N values

Soil nitrogen isotope ratios (i.e. bulk soil δ^15^N values) were measured every two years from 50 mg of dried soil (after grinding with a ball-mill) with an IRMS (Delta C prototype IRMS, Finnigan MAT)^73^.

##### S1.1.32. Soil NH_4_ and soil NO_3_

Each autumn from 2002 to 2008, five soil cores (diameter 0.01 m) were taken at a depth of 0 to 0.15 m of the mineral soil from each of the experimental plots and pooled. As an estimate of plant□available N, NO_3_□N and NH_4_□N concentrations were determined by extraction of soil samples with 1 M KCl solution^71^. Nitrate□N and NH_4_□N concentrations were measured in the soil extract with a Continuous Flow Analyzer (CFA, 2003–2005: Skalar, Breda, Netherlands; 2006–2008: AutoAnalyzer, Seal, Burgess Hill, United Kingdom).

##### S1.1.33. Nitrate leaching

Nitrate leaching was calculated by multiplying soil NO3 concentrations (see S1.1.32) with downward fluxes of soil water (0-30 cm depth) (S1.1.27).

##### S1.1.34. Soil Phosphate

Concentrations of soil phosphate were determined in soil solution, which was collected every two weeks (cumulative sample) between 2003 and 2007, 2009, 2011 and 2012 using suction plates with permanent vacuum at 30cm soil depth. Soil solution samples were then analysed photometrically with Continuous Flow Analysis (CFA; see 1.1.32). From these biweekly measurements, an annual average was calculated for each plot.

#### S1.2. Trait measurements

**Table S1.2:** Overview of traits

| Trait | Unit | Description |
| --- | --- | --- |
| shoot:root ratio | $\text{g g}^{-1}$ | Shoot mass per root mass |
| shoot:root N ratio | unitless | Leaf nitrogen uptake / root nitrogen uptake |
| plant height | cm | Standing height of the shoot |
| leaf biomass production rate | $\text{g day}^{-1}$ | Maximum daily leaf dry mass production |
| total leaf area | $\text{cm}^2$ | Total area of all leaves of plant |
| leaf area | $\text{mm}^2$ | Average area of a single leaf |
| leaf thickness | mm | Leaf thickness |
| specific leaf area | $\text{mm}^2 \text{g}^{-1}$ | Fresh leaf area per leaf dry mass |
| leaf specific density | $\text{g cm}^{-3}$ | Leaf dry weight per leaf fresh volume |
| leaf area ratio | $\text{cm}^2 \text{g}^{-1}$ | Leaf area per shoot mass |
| leaf form coefficient | $\text{mm}^2 \text{mm}$ | Leaf area divided by leaf perimeter |
| leaf dry matter content | $\text{g g}^{-1}$ | Leaf dry weight per leaf fresh weight |
| leaf C content | % | Leaf carbon content |
| leaf N content | % | Leaf nitrogen Content |
| leaf conductance | $\mu\text{M s}^{-1} \text{A}^{-1}$ | Stomatal conductance per leaf area |
| leaf toughness | N | Leaf resistance to penetration |
| stem diameter | mm | Diameter of stem |
| stem specific density | $\text{g cm}^{-3}$ | Stem dry weight per stem fresh volume |
| erectness | $\text{cm cm}^{-1}$ | Stretched height per standing height |
| biomass fraction inflorescence | $\text{mg mg}^{-1}$ | Inflorescence:shoot biomass fraction |
| inflorescences per shoot | nr | Number of inflorescences per shoot |
| duration flowering | ordinal | Duration of flowering period |
| seeds projected area | $\text{mm}^2$ | Total area of individual seed |
| nr seedlings | nr | Number of plant seedlings within subplot |
| seed weight | g | Weight of 1000 seeds |
| seed width length ratio | $\text{mm mm}^{-1}$ | Ratio of seed width to seed length |
| seed dry matter content | $\text{g g}^{-1}$ | Seed dry weight per seed fresh weight |
| root area | $\text{cm}^2$ | Root area |
| rooting depth | ordinal | Depth of the root system |
| root area distribution | unitless | Evenness of vertical root area distribution |
| specific root area | $\text{cm}^2 \text{g}^{-1}$ | Root surface area per root mass |
| specific root length | $\text{cm g}^{-1}$ | Root length per root mass |
| root tissue density | $\text{g cm}^{-3}$ | Root dry weight per root volume |
| root nitrogen uptake | $\text{mg day}^{-1}$ | Nitrogen uptake into roots |
| root CN ratio | unitless | Root total carbon:nitrogen content |
| root P content | ‰ | P content per root dry biomass |
| root K content | ‰ | K content per root dry biomass |
| root S content | ‰ | S content per root dry biomass |
| root Ca content | ‰ | Ca content per root dry biomass |
| root Na content | ‰ | Na content per root dry biomass |
| nutrient uptake efficiency | $\text{mg g}^{-1}$ | Root nitrogen uptake:root biomass |

Most of the functional traits listed in Table S1.2 (except for the seed traits and biomass fraction of inflorescences, number of inflorescences per shoot and number of seedlings) were measured in mesocosms. To this end, we obtained seeds of all 60 plant species used in the Jena Biodiversity Experiment from a seed supplier (Rieger Hoffmann GmbH, Blaufelden-Raboldshausen, Germany and Saaten Zeller e.K., Riedern, Germany). In April 2011 and 2012 we germinated the seeds in petri dishes and we planted seedlings of 1-3 weeks old into mesocosms, with for each species five replicates. Seedlings that dead within 4 weeks after transplanting were replaced. Mesocosms were made of PVC pipes (height = 60 cm, diameter = 15 cm). Mesocosms were placed outside in the Botanical Garden of Leipzig (Germany), in randomized blocks. Traits were measured after 12 weeks. For more details of the mesocosm design, we refer to Schroeder-Georgi *et al.*^6^.

For detailed methods on the trait measurements of shoot:root ratio, plant height, leaf biomass production rate, total leaf area, leaf area, leaf thickness, specific leaf area, leaf specific density, leaf area ratio, leaf dry matter content, leaf C content, leaf N content, leaf conductance, leaf toughness, stem specific density, erectness, root area distribution, specific root area, specific root length, root tissue density, root nitrogen uptake, root C:N ratio, we refer to Schroeder-Georgi *et al.*^6^. Shoot:root N ratio was calculated as the leaf nitrogen uptake divided by the root nitrogen uptake, based on measurements of Schroeder-Georgi *et al.*^6^. Leaf form coefficient was calculated as the leaf area (see above) divided by the leaf perimeter. Leaf perimeter was measured on the same picture from samples as leaf area, using the software WinFolia (Regent Instruments Inc., Canada). Stem diameter was measured on the same stems as those used for stem specific density^6^ and defined as the diameter of a stem in mm. Nitrogen uptake efficiency was calculated as the root nitrogen uptake divided by the root dry biomass (measurements from Schroeder-Georgi *et al.*^6^). Root area was based on the root area measurements of Schroeder-Georgi *et al.*^6^. Duration of flowering was defined as the duration of the flowering period, and expressed using an ordinal scale: 1 (1 month), 2 (2 months), 3 (3 months) and 4 (more than three months). Root element contents (P, K, S, Ca, Na) were analyzed using a subsample of dried fine root material of each mesocosm. A microwave digestion system (Berghof Speedwave SW-2) was used to digest 0.2 g ground material for 50 min at 190° using 8ml HNO3, 3ml H2O2. The method was tested using standard reference material. Samples were analyzed using ICP-OES (Spectro Acros, Spectro Analytical Instrument). Seed traits were measured on a subsample of the seeds purchased for the mesocosm experiment (see above). Seeds were cleaned from all attached tissue (e.g. bracts from grass spikelets), placed in batches of 30 - 200 well apart in glass petri dishes and scanned using a flatbad scanner (resolution 800 dpi) and analyzed using WinSeedle (Reg. 2009a, Regent Instruments Inc., Canada). WinSeedle output provided data on seed length, seed width and seed projected area for individual seeds from each image. Seed projected area and seed width to length ratio were calculated as mean over individual seed measures per species. Seed batches were weighed fresh, dried (70°, 48 h), and weight again to calculate seed dry matter content as dry weight per fresh weight for the total seed batch and the weight of 1000 seeds per species using the seed number measured with WinSeedle and seed dry weight. Data on duration of flowering was obtained from Roscher et al. 2014^20^. Rooting depth was also obtained from Roscher et al. 2014^20^. It was measured on an ordinal scale: 1 (up to 20 cm), 2 (up to 40 cm), 3 (up to 60 cm), 4 (up to 100 cm) and 5 (> 100 cm). Biomass fraction of inflorescence (mg_inflorescence_ mg^-1^ _hoot_) and number of inflorescences per shoot were recorded in the small-area monocultures of the field experiment (between 2006 and 2009) or in a low-diversity mixture for three species not abundant enough in the monocultures. Five to seven shoot per species were sampled. In the laboratory, the number of inflorescences per shoot was counted. Afterwards shoots were separated into compartments (stems, leaves and reproductive parts), the compartments were dried (48 h, 70°C) and weighed. The mass of reproductive parts was divided by summed biomass of all compartments per shoot to derive inflorescence mass fraction^74^. The number of seedlings (i.e. plant individuals with cotyledons) was counted in all small-area monocultures three times (April, July, October) in 2007 to account for species-specific differences of seedling emergence. Three quadrats of 0.3 × 0.3 m size per subplot were randomly placed for each census. Total numbers of emerged seedlings per m^2^ were calculated for each monoculture based on pooled data from all census dates^74^.

**Table S1.3.**
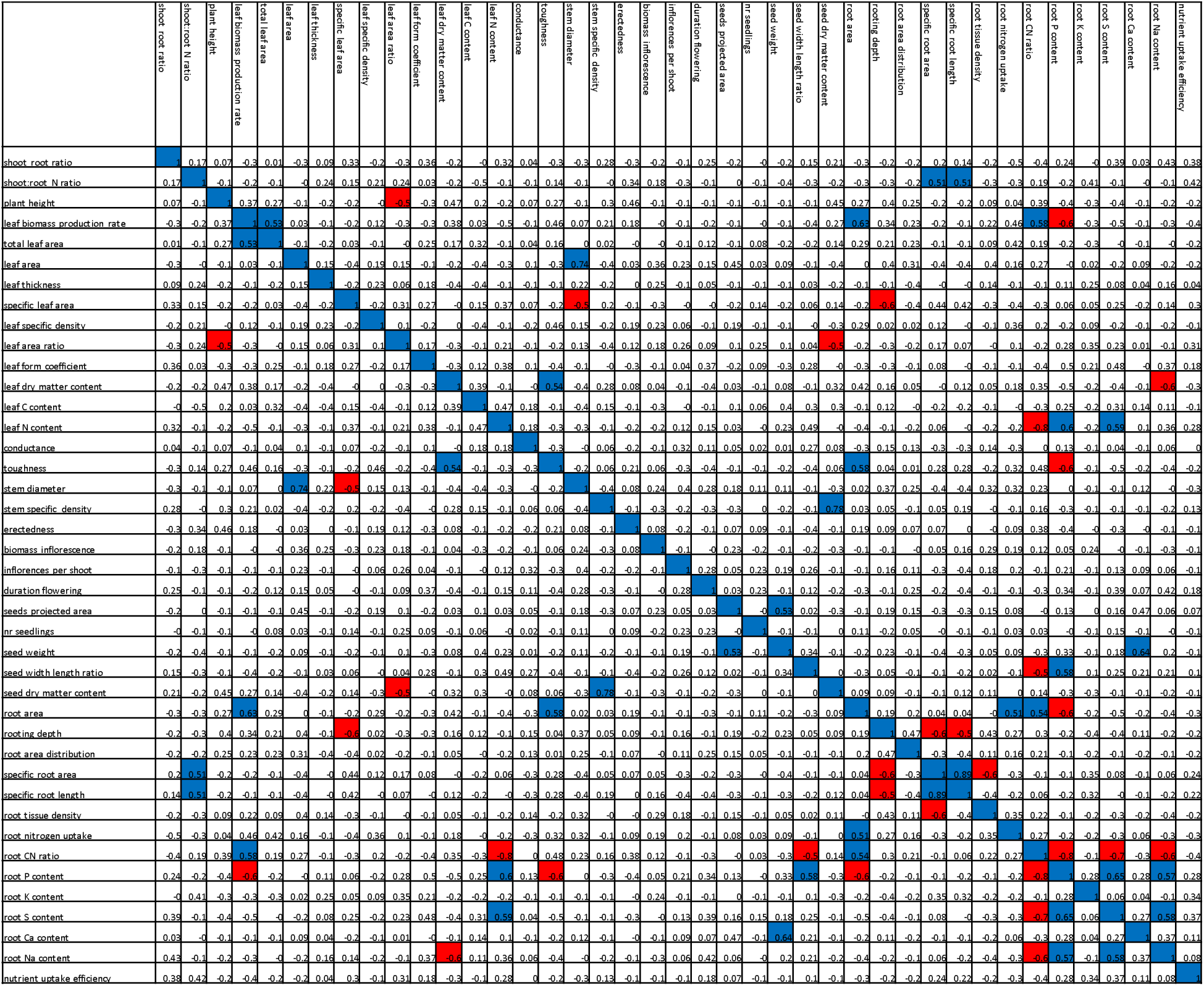
Pearson correlation coefficients between traits.

### S2. SUPPLEMENTARY RESULTS

#### S2.2. Overview of final model outcomes

On average, each trait significantly affected 4.9 out of the 42 ecosystem functions in the final models, and each ecosystem function was driven by 4.8 different traits. However, traits varied in the identity and number of ecosystem functions they drove, and vice versa, ecosystem functions varied in the identity and number of traits by which they were driven. Table S.2.1 gives an overview of which traits (their functional identity and/or their functional diversity) were significantly driving which functions in final models. Average marginal R^2^ values of models were 0.127. This was slightly lower (0.121) when FI and FD metrics based on presence-absence data (instead of abundance data) were used as predictors.

**Table S2.1.**
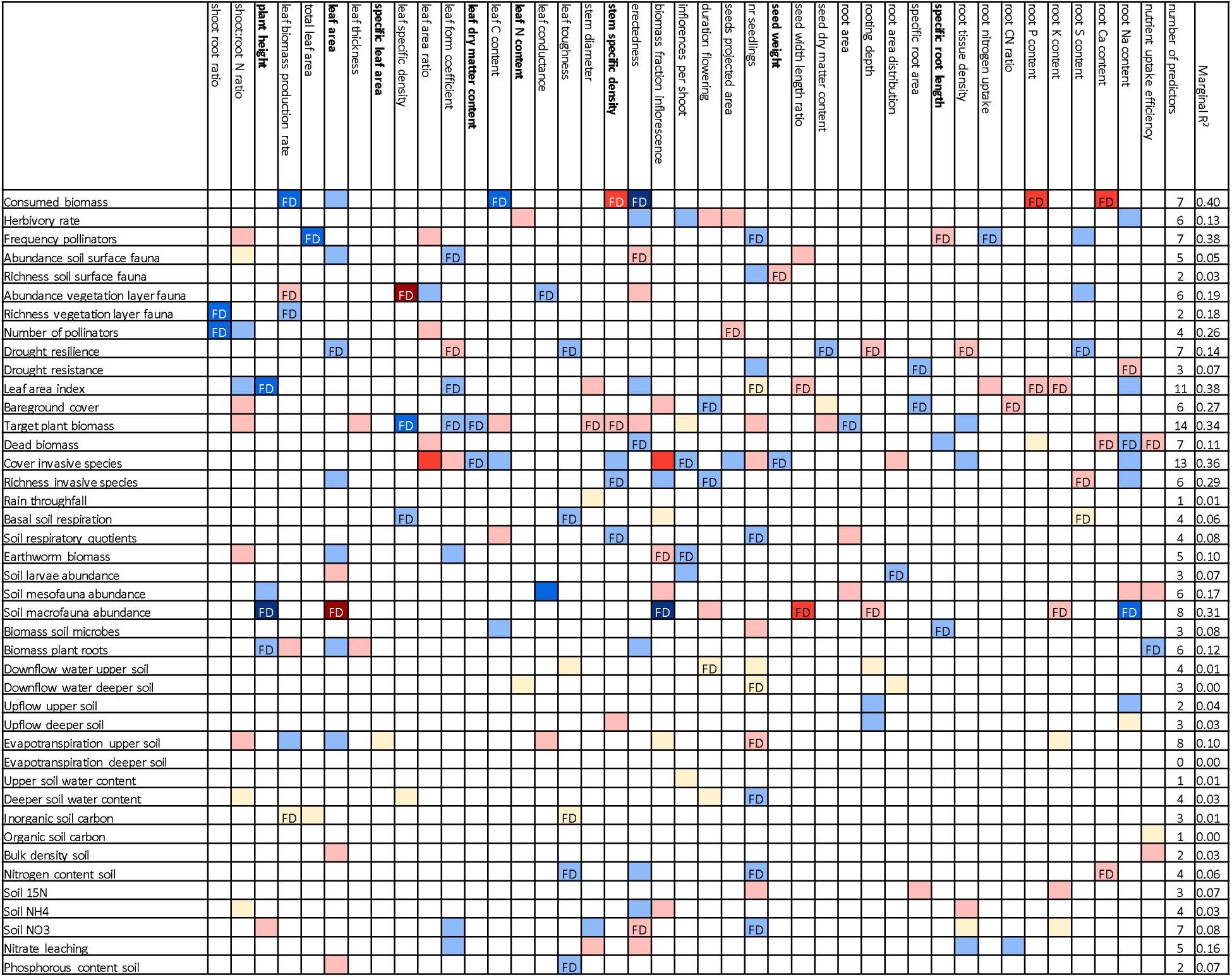
Ecosystem functions and their relationships with plant traits. Colored squares indicate whether the Functional Diversity and/or Community Weighted Mean of a given trait was present in the final model explaining the corresponding ecosystem function, and whether the effect was strongly negative (dark red, *r* < −0.5), moderately negative (normal red, −0.5 ≤ *r* < −0.3), weakly negative (light red, −0.3 ≤ *r* < −0.1), neutral (yellowish, −0.1 ≤ *r* < 0.1), weakly positive (light blue, 0.1 ≤ *r* < 0.3), moderately positive (normal blue, 0.3 ≤ *r* < 0.5) or strongly positive (dark blue, *r* < 0.5). When the Functional Diversity of the trait was the strongest predictor, FD is written in the cell; in all other cases, Functional Identity of the trait was the strongest predictor. The ecosystem functions analyzed in over 10% of the papers included in the mini-review are shown in bold. At the end of each row, a number is given indicating how many traits were significantly related to the corresponding ecosystem function. Similarly, at the bottom of each column, a number is given indicating how ecosystem functions were significantly related to the corresponding trait.

